# Sensitive and scalable metagenomic classification using spaced metamers, reduced alphabets, and syncmers

**DOI:** 10.64898/2026.03.13.711249

**Authors:** Jaebeom Kim, Martin Steinegger

## Abstract

Accurate taxonomic classification of metagenomic sequencing data is essential for identifying the diverse organisms present in environmental and clinical samples. To address this, we previously developed Metabuli, an alignment-free classifier that bridges the gap between nucleotide-level resolution and protein-level sensitivity. Central to this approach is the ‘metamer’, a novel joint DNA and amino acid *k*-mer concept introduced by Metabuli to classify taxa across varying levels of database representation. In this study, we significantly optimized its core metamer search by incorporating advanced techniques. By introducing spaced metamers and reduced amino acid alphabets to boost sensitivity, we improved both precision and recall by 4.7 and 2.8 percentage points in a species exclusion test. Furthermore, integrating syncmers with spaced metamers halved the reference database size and doubled the classification speed; despite a slight decrease in recall, this configuration continued to outperform state-of-the-art alignment-free tools.

## Introduction

Metagenomic studies analyze uncharacterized DNA sequences recovered from diverse environments to profile the taxonomic composition of complex microbial communities. Assembly-free approaches can be broadly categorized into two types: sample-centric and read-centric analyses (1; 2). The former extracts global features from the entire sample such as *k*-mer sketches and compares them to reference databases, enabling fast species containment and composition estimation (3; 4). The latter approach identifies the taxonomic origin of each individual read (2). While read-level classification is computationally heavier and inherently slower, it can easily aggregate into sample-level compositions while providing critical granularity for downstream analyses. Furthermore, because read-centric approaches evaluate every sequence rather than relying on global sketches, they are less susceptible to low sequencing coverage and are superior at detecting rare or novel species. In this work, we focus on the latter approach.

Read-centric classifiers search homologous regions between query and reference sequences. Alignment-based methods can achieve high sensitivity at a significant computational cost (5). Exact *k*-mer matching algorithms provide ultra-fast classification, though their fixed-length requirements suffer when divergent sequences are encountered (6). It is partially mitigated by suffix-based approaches, which utilize data structures like the FM-index to identify variable-length maximal exact matches (7; 8; 9). More recently, approximate mapping and pseudoalignment algorithms have emerged, utilizing seed chaining to estimate read origins without the overhead of dynamic programming.

While often categorized separately, exact *k*-mer matching serves as the primary filtering engine across these algorithmic approaches. Consequently, numerous techniques have been developed to optimize its sensitivity and efficiency. To enhance sensitivity, for example, spaced *k*-mers incorporate internal ‘joker’ positions to tolerate mismatches at specific sites (10). For amino acid *k*-mers, reduced alphabets group physicochemically similar amino acids into representative classes, enabling algorithms to overlook conservative substitutions (11; 12). To address efficiency, sub-sampling strategies such as minimizers (13) and syncmers (14) are employed to select a representative subset of k-mers from a sequence (15). By only indexing and querying the subset, these methods drastically reduce memory requirements and computational overhead while maintaining sufficient sequence coverage, decreasing the storage burden as well.

In our previous work, we integrated DNA- and amino acid-based *k*-mer matching to combine nucleotide-level resolution with protein-level sensitivity (16). To achieve this, we introduced the metamer, a novel *k*-mer data structure that simultaneously stores both amino acid and DNA information. For compact encoding, the metamer exploits the redundancy of the genetic code; because there are at most six synonymous codons per amino acid, the original DNA sequence can be retained using only 3 bits per codon. We implemented this approach in the classifier Metabuli, which identifies exact matches at the amino acid level to maximize sensitivity, and then utilizes the underlying DNA information to differentially weight those matches. Metabuli consistently achieved top-tier performance across benchmarks that demand either nucleotide-level resolution or protein-level sensitivity, demonstrating the successful integration of both strengths. Additionally, we developed Metabuli App (17), a cross-platform desktop application that pairs Metabuli with integrated read quality control and interactive visualizations within an intuitive graphical interface.

In this study, we further enhanced Metabuli’s performance by integrating advanced *k*-mer optimization techniques into the core metamer framework. First, we developed a flexible metamer encoding scheme to facilitate seamless fusion of new techniques. To boost sensitivity, we introduced reduced alphabets and spaced metamers (Figure 1), and for efficiency, we implemented closed syncmer sampling (14). We chose syncmers over minimizers for their context-independence: the selection of a syncmer depends entirely on the *k*-mer itself, remaining unaffected by neighboring mutations. This ensures that if a *k*-mer is selected in a query sequence, it will deterministically be selected in the reference as well. Moreover, closed syncmers provide a distance guarantee, enabling predictable *k*-mer chaining.

**Figure 1.**
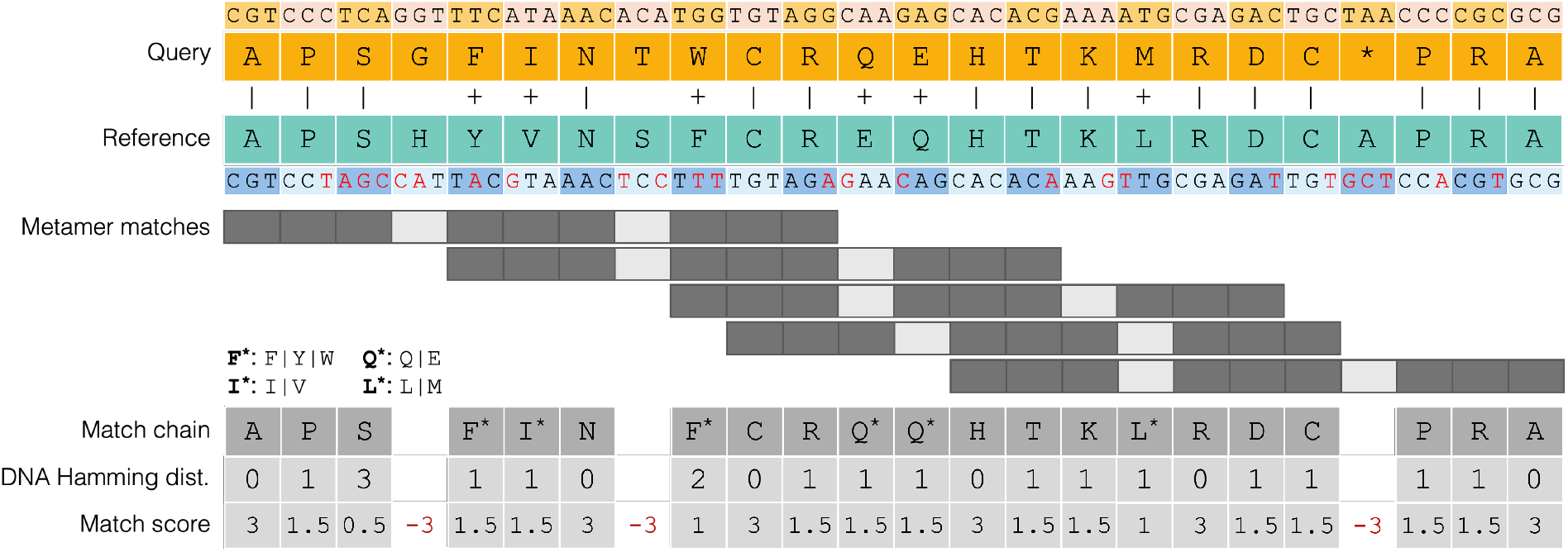
Robust homology detection via spaced metamer chaining. Alignment of diverged query and reference sequences demonstrating the combined use of reduced amino acid alphabets and spaced seeds. Identical residues are indicated by (|). Conservative substitutions (+) are tolerated through exact matching using reduced amino acid alphabets (e.g., F* grouping F, Y, and W). The spaced metamer’s joker positions enables matching across amino acid mismatches at positions 4, 8, and 21. Overlapping metamers are merged into a single match chain, which is subsequently scored based on the underlying DNA Hamming distance (mutations are red in Reference) and spaces to provide a nucleotide-aware match score.

We assessed the individual and combined impacts of these techniques on classification resolution and sensitivity using both species inclusion and exclusion benchmarks. Finally, we compared the enhanced Metabuli configurations against state-of-the-art alignment-free tools across a wide range of taxonomic ranks.

## Methods - Implementations

While the fundamental framework for database construction and read classification remains consistent with the original Metabuli architecture, we have introduced several significant methodological refinements to enhance sensitivity and statistical rigor. This section details these integrations.

### Flexible metamer encoding and introduction of a reduced amino acid alphabet

A metamer is a *k*-mer data structure that simultaneously encodes an amino acid sequence of *k* residues and their originating codons. Previously, Metabuli only supported the standard translation table for this metamer encoding.

Users can now define a custom metamer encoding via a flexible translation table, provided the total *k*-mer fits within a 64-bit integer limit. Specifically, the table must satisfy the constraint *k* × (*b*_*aa*_ + *b*_*c*_) ≤ 64, where *b*_*aa*_ = ⌈log_2_ |*A*|⌉ represents the bits required to encode the amino acid alphabet of size |*A*| including one stop codon, and *b*_*c*_ = ⌈log_2_(*C*_*max*_)⌉ represents the bits needed to encode the maximum number of synonymous codons mapping to a single amino acid (*C*_*max*_).

This flexibility enables the use of reduced amino acid alphabets for metamer encoding. We opted for a reduced alphabet introduced by Li et al. (11), which groups specific amino acids together: (F, Y, W), (M, L), (I, V), and (Q, E). This reduction yields an alphabet size of |*A*| = 16, requiring *b*_*aa*_ = 4 and *b*_*c*_ = 3. This allowed us to encode a 9-mer instead of Metabuli’s default 8-mer, which compensates for the reduced specificity of the compressed alphabet.

### Reference and query metamer extraction

Metamer extraction begins by translating DNA sequences according to a specified translation table. As established in the Metabuli framework (16), reference reading frames are predicted using Prodigal, while query sequences undergo a six-frame translation. T Metamers are then extracted from the amino acid sequences and their original codons

### Changes in metamer encoding scheme

Previously, Metabuli encoded amino acid 8-mer into a 36-bit binary vector by treating the sequence as a base-21 integer, calculated as 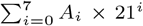 where *A*_*i*_ ∈ {0, 1, …, 20} represents the integer assignment for the 20 standard amino acids and the stop codon at position *i*. Storing the corresponding codon information required an additional 24 bits (*b*_*c*_ = 3 for the standard translation table), resulting in a total of 60 bits per metamer. While highly compact, this encoding scheme introduced significant overhead: it required calculating the polynomial for every metamer extraction and made retrieving the original amino acid and DNA strings from the k-mer computationally expensive.

To overcome this computational overhead, the updated metamer encoding now utilizes a direct bit-packing strategy. Within a 64-bit C++ integer, the data is packed such that the *k* × *b*_*c*_ least significant (rightmost) bits encode the specific synonymous codon choices, while the adjacent *k* × *b*_*aa*_ bits to the left encode the amino acid sequence. Within each of these distinct blocks, the sequence is encoded directionally: data for the leading amino acid is placed in the most significant (leftmost) bits of its respective block, sequentially mapping the *k*-mer down to the rightmost bits. This layout allows for rapid sequence retrieval through simple bit-shifting operations.

This layout enables highly efficient *k*-mer extraction using a sliding window and simple bitwise operations. Instead of recalculating the entire *k*-mer, running bit-vectors for the amino acid and codon sequences are incrementally updated for each new triplet via left-shifts (*b*_*aa*_ and *b*_*c*_) and bitwise OR operations. To assemble the final metamer, masks trim the vectors to *k* residues, and the amino acid block is shifted left by *k* × *b*_*c*_ bits to merge with the codon block. The sliding window automatically resets upon encountering ambiguous bases (e.g., ‘N’) to ensure only valid, contiguous metamers are extracted.

### Syncmer-based sub-sampling

Building upon this optimized bitwise architecture, we implemented an efficient closed syncmer extraction algorithm utilizing a double-ended queue (deque), inspired by Kraken2’s minimizer selection algorithm (6). To identify closed syncmers—defined by the presence of the minimum *s*-mer at start or end positions within the *k*-mer—a sliding window of *s*-mers is continuously updated via bitwise shifts of the translated amino acid sequence. The deque maintains the minimum *s*-mer within the current window by discarding larger or out-of-bound values. When the minimum *s*-mer aligns with the required start or end anchors, the sequence is validated as a closed syncmer. To maximize speed during metamer assembly, the algorithm avoids full recalculation; it incrementally shifts and appends only the new amino acid and codon bits corresponding to the positional offset from the previously identified syncmer.

To facilitate exact amino acid matching, syncmer selection criteria were applied solely to the amino acid sequences of the metamers. Consequently, all metamers sharing identical amino acid sequences are co-selected or co-rejected.

### Spaced metamer and syncmer extraction

We extended the metamer framework to support spaced seeds, defined by a mask of span *w* and weight *k* (*w > k*). The process involves two stages: *w*-mer extraction followed by *k*-mer compaction. The *w*-mer is first extracted identically to a contiguous *k*-mer. Next, the sequence is compacted into a dense representation by filtering out the masked positions and tightly packing only the active residues into a 64-bit integer. To optimize this compaction, our implementation utilizes hardware-level intrinsics—specifically counting trailing zeros (`--builtin_ctz`)—to rapidly isolate and pack bits from the masked positions.

For spaced syncmer extraction, the syncmer condition can be defined by evaluating *s*-mers within either the original *w*-mer span or the condensed *k*-mer sequence. While the former allows for an efficient deque-based implementation, it compromises the context-independence of the syncmer; residues at inactive mask positions—which are not included in the final encoding—would influence the selection. Therefore, we define the syncmer condition using the condensed *k*-mer to ensure that selection is determined solely by the residues being encoded.

### Metamer chaining for syncmer and spaced metamer

To compensate for the low specificity of default 8-mers, Metabuli requires individual metamer matches to form a contiguous chain spanning at least eleven amino acids. Our previous work introduced an efficient, single-pass algorithm to simultaneously generate and score these chains (17), linking adjacent matches shifted by one amino acid by verifying the exact identity of their 21 overlapping DNA bases.

We have now extended this algorithm to support both syncmers and spaced metamers. Because syncmers are sub-sampled, matches shifted by a single amino acid are infrequent. However, because closed syncmers offer a bounded window guarantee of (*k* − *s*), the algorithm is now extended to chain matches shifted by up to *k* − *s* positions, requiring exact identity across their 24 − 3(*k* − *s*) overlapping DNA bases.

For spaced metamers, matches shifted by a single amino acid are not viable when mutations occur at the joker positions; requiring strictly adjacent matches would negate the mismatch tolerance provided by the spaces. Consequently, the algorithm permits larger shifts to bypass these jokers. These permissible shifts are further extended when syncmer sampling is applied together; for instance, the syncmer settings in Table 1 allow for shifts of up to eight amino acids. Because of the joker positions, matches shifted by *n* positions may share fewer than *k*−*n* residues, and the specific locations of these shared residues depend on how the active positions of the two shifted masks align. To dynamically compute these variable overlaps, we utilize hardware intrinsics and bitwise operations to efficiently define and extract the shared residues between two spaced metamers shifted by *n* amino acids. The resulting metamer chain is scored by adding 3, 1.5, 1, or 0.5 for (reduced) amino acid matches with DNA Hamming distances of 0 to 3, respectively, and applying a penalty of 3 for spaces (Figure 1).

**Table 1.** Summary of the exclusion test set. Entries represent genome counts (taxon counts). The columns detail the dataset prior to exclusion (Before), the held-out query set (Exclusion), and the resulting dataset (After)

| Rank | Before | Exclusion | After |
| --- | --- | --- | --- |
| Family | 53,941 (1,125) | 337 (56) | 53,604 (1,069) |
| Genus | 53,604 (3,480) | 5,468 (149) | 48,136 (3,331) |
| Species | 48,136 (12,922) | 3,811 (478) | 44,325 (12,444) |
| Subspecies | 44,325 (44,325) | 1,020 (1,020) | 43,305 (43,305) |

### Selection of spacing mask

For standard codon-based 8-mers, we adhered to Metabuli’s established rule requiring a chain to span at least eleven amino acids. Because adding three spaces would cause the minimal chain of two adjacent matches to overextend to twelve amino acids, we utilized two spaces—the maximum allowable limit. Conversely, our 9-mer implementation relies on a reduced amino acid alphabet, which inherently lowers sequence specificity. To compensate, we added two spaces to the 9-mer ensures that the minimal chain spans twelve residues.

Incorporating two ‘joker’ (*J*) positions partitions the mask into three active contiguous blocks: *X* − *J*_1_ − *Y* − *J*_2_ − *Z*. To bypass mismatches occurring at both jokers, a subsequent match must align its *X* block with either the *Y* or *Z* block of the preceding match. The first case yields a total span of *L*(*X*) + 2*L*(*Y*) + *L*(*Z*), whereas the second case appends an additional max(*L*(*X*), *L*(*Z*)) penalty term to this span. If *L*(*X*) *> L*(*Y*), only the second case is viable; this physically increases the number of residues required to form a valid chain, making chaining overly stringent. Therefore, we enforce the constraint *L*(*X*) ≤ *L*(*Y*).

If *L*(*Y*) *< L*(*Z*), the succeeding match’s *J*_2_ falls within the preceding exact-match region (*Z*), wasting its mismatch tolerance. Thus, we require *L*(*Z*) ≤ *L*(*Y*). Minimizing *L*(*X*)+ 2*L*(*Y*) + *L*(*Z*) subject to *L*(*X*), *L*(*Z*) ≤ *L*(*Y*) yields the optimal 9-mer configuration: *L*(*X*) = *L*(*Y*) = *L*(*Z*) = 3. For 8-mers, this results in *L*(*Y*) = 3, with *L*(*X*) and *L*(*Z*) being 2 and 3. Consequently, we implemented the mask pattern `11101110111` for 9-mers utilizing the reduced alphabet, and `1110111011` for standard codon-based 8-mers.

### E-value calculation

To assess the statistical significance of exact metamer chains, we implemented a composition-aware E-value. Because traditional alignment statistics are overly stringent for discrete exact matches, we model the log-transformed E-value as:

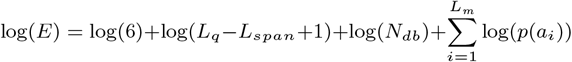

where the factor of 6 accounts for the six-frame translation of the query, *L*_*q*_ is the query length in amino acids, *L*_*span*_ is the match span, M is the number of exact matches, and *N*_*db*_ is the database size. *N*_*db*_ was calculated by dividing the total length of reference genomes by 3. Here, *p*(*a*_*i*_) is the background probability of the *i*-th matched character. This approach naturally penalizes promiscuous random matches while log-space computation ensures stability for long chains. An E-value-based filtering option is also provided, with a default threshold of 1.0.

### Changed tie criterion for LCA calculation

Previously, alternative matches were considered a tie if their score exceeded 0.95 × the highest score. For a query length of 300 with a perfect normalized top score of 1.0, this allowed an absolute score difference of 15, which could encompass up to 10 DNA mismatches. Because this reduced the number of species-specific classifications, we increased the tie ratio to 0.99 for perfect matches. However, applying this stringent criterion uniformly to lower-scoring matches resulted in overconfident classifications. To balance this, we implemented a dynamic tie ratio defined as 0.99 − 0.09 ×(1 − *S*_*best*_), where *S*_*best*_ is the normalized best score.

## Methods - Benchmark details

### Generating a synthetic benchmark dataset

We designed a benchmark dataset to evaluate classification improvements according to the query-to-reference distance, consisting of a single reference genome set and multiple query genome sets. Each query genome set is either for inclusion or exclusion tests at different ranks. An inclusion test at a certain rank evaluates discriminating resolution at the rank, while an exclusion test assesses homology detection sensitivity at the rank. For example, a query set for the species-inclusion test contains genomes of species if both the target species and its sibling species are included in the reference set. In the case of the species-exclusion test, a query set consists of genomes of species where the target species is absent, but its sibling species are included in the reference set.

We first downloaded assemblies at the complete-genome or chromosome level with CheckM2 completeness*>*90 and contamination*<*5 from GTDB release 226 (18), which resulted in a pool of 53,941 genomes across 12,444 species.

Query-set generation begins with the family-exclusion test. From orders containing multiple families in the genome pool, it randomly selects one-third. For each selected order, it randomly chooses one family and excludes all genomes belonging to it from the genome pool. From each excluded family, it then selects one genome to simulate query reads for the family-exclusion test. Using the reduced genome pool, the same procedure is applied at the genus level for genus-exclusion test. This procedure is subsequently repeated for the species- and subspecies-exclusion tests.

After all exclusions, the pool was reduced to 43,305 genomes, which comprised the single reference genome set (Table 1). To construct databases for the protein-based classifiers, we downloaded the corresponding annotated protein FASTA files (`*_protein.faa.gz`) when available from NCBI. For assemblies lacking pre-annotated proteins, we utilized Pyrodigal (19) to predict the protein sequences. To construct the query set for the subspecies-inclusion test, a random eighth of the species with multiple genomes in the reference set were selected (329 species), and two genomes were randomly chosen per selected species. For the species-inclusion test, a random quarter of the genera with multiple species in the reference set were selected (309 genera), and from each selected genus, two genomes representing different species were randomly chosen. The generation of these inclusion and exclusion test sets is fully automated via the `maketestsets` and `makeInclusionTestQueries` modules within Metabuli. For each query set, paired-end reads were simulated from its genomes using Mason2 (v2.0.9) (20). Simulation parameters followed Kim et al. (16); only the coverage varied between tests.

### Performance metrics

Performance was evaluated at the most specific taxonomic rank allowed by the benchmark constraints (e.g., genus-level evaluation for species-exclusion tests). At a given target rank, True Positives (TPs) were defined as assignments consistent with the ground-truth lineage at that rank or any more specific sub-clade. To penalize low-resolution assignments, unclassified reads and those resolved only to ranks above the target were categorized as False Negatives (FNs). Any assignment to an incorrect taxonomic lineage was recorded as a False Positive (FP). Based on these categories, we calculated precision, recall, and the F1-score as follows: Precision 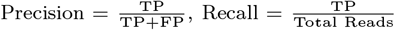 and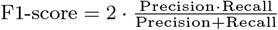

### Benchmarked software

Kraken 2 (v.2.17.1) (6) was run using default settings, with the addition of the `-–protein` parameter to construct the database for translated searches (Kraken2X). Kaiju (v1.10.1) (9) and Centrifuger (v1.1.0) (7) were executed with default parameters.

## Results and Discussion

### Reduced amino acid alphabets and spacing improves sensitivity

We evaluated the effect of reduced amino acid alphabets and spaced k-mers on classification resolution and sensitivity across six configurations of Metabuli v1.2, using v1.1.1 as a baseline (Table 2). E-value filtering was disabled for compatibility with v1.1.1. Although both v1.1.1 and the first configuration of v1.2 use standard alphabets and contiguous k-mers, their performance differed primarily due to the updated LCA tie criterion.

**Table 2.** Performance Comparison of different *k*-mer technique combinations. *k* denotes *k*-mer length; *L*_*min*_ denotes the minimum number of covered residues; *j* denotes the number of joker positions. E-value-based filtering was disabled.

| Alphabet | *k* | *L<sub>min</sub>* | *j* | Pre. | Recall | F1 |
| --- | --- | --- | --- | --- | --- | --- |
| Species exclusion test |  |  |  |  |  |  |
| v1.1.1 | 8 | 11 | 0 | 0.735 | 0.595 | 0.658 |
| Standard | 8 | 11 | 0 | 0.758 | 0.590 | 0.663 |
| Reduced | 8 | 11 | 0 | 0.710 | 0.600 | 0.650 |
| Reduced | 9 | 11 | 0 | 0.742 | 0.611 | 0.670 |
| Reduced | 9 | 12 | 0 | 0.767 | 0.599 | 0.673 |
| Standard | 8 | 11 | 2 | 0.764 | 0.610 | 0.678 |
| Reduced | 9 | 12 | 2 | 0.768 | 0.629 | 0.691 |
| Species inclusion test |  |  |  |  |  |  |
| v1.1.1 | 8 | 11 | 0 | 0.9997 | 0.9791 | 0.9893 |
| Standard | 8 | 11 | 0 | 0.9993 | 0.9863 | 0.9928 |
| Reduced | 8 | 11 | 0 | 0.9993 | 0.9885 | 0.9939 |
| Reduced | 9 | 11 | 0 | 0.9992 | 0.9862 | 0.9926 |
| Reduced | 9 | 12 | 0 | 0.9992 | 0.9864 | 0.9928 |
| Standard | 8 | 11 | 2 | 0.9992 | 0.9872 | 0.9932 |
| Reduced | 9 | 12 | 2 | 0.9993 | 0.9873 | 0.9932 |

The species exclusion test requires high sensitivity to detect homology between sibling species for accurate genus-level classification. While adopting reduced alphabets increased recall, their lower specificity caused a drop in precision, lowering the overall F1-score. Increasing the *k*-mer length to 9 successfully recovered both metrics. Subsequently, raising the minimum covered residues from 11 to 12 yielded precision and recall rates strictly higher than the standard 8-mer baseline. Finally, integrating a spaced mask provided substantial gains, improving precision and recall by 3.3 and 3.4 percentage points, respectively, over Metabuli v1.1.1. Figure 1 demonstrates how a match is formed under this final configuration.

Conversely, in the species inclusion test, we did not observe significant changes in species-level resolution. This occurs because the metamers still retain the essential DNA information needed to distinguish closely related species. As the tie criterion for LCA calculation became more stringent for high-scoring matches in the new version of Metabuli, species-level recall increased alongside a drop in precision, resulting in a higher overall F1 score compared to v1.1.1. We utilized the last configuration for subsequent analyses with maximum E-value set as 1.0, maintaining the v1.1.1 as a baseline.

### Closed syncmer improves the classification scalability

The invariance of closed syncmer selection to adjacent mutations facilitates exact amino acid matches, while its distance guarantee allows for the chaining of overlapping metamers. Given query and reference *k*-mer sets of sizes *A* and *B* with *X* shared *k*-mers, independent random subsampling at a 50% rate would reduce the shared *k*-mers to *X/*4, dropping the match density to *X/*(2(*A* + *B*)). Syncmer selection, however, is deterministic; it retains *X/*2 shared *k*-mers, preserving the original density of *X/*(*A* + *B*) and thereby promoting robust *k*-mer matching.

By varying *s*-mer length, we assessed the database size, classification speed, and accuracy (Table 3). Our implementation of closed syncmer selection on spaced metamers successfully achieved the theoretical compression rate of (*k* − *s* + 1)*/*2. Furthermore, classification speed was accelerated proportionally to this compression rate.

**Table 3.** Effect of using syncmers. *s, c*, and *d* denote *s*-mer length, expected compression rate, and match density, respectively. The final configuration of Table 2 is used.

| $s$ | $c$ | DB size<br>(GB) | Speed<br>(reads/s) | Pre. | Recall | F1 |
| --- | --- | --- | --- | --- | --- | --- |
| Species exclusion test |  |  |  |  |  |  |
| - | 1 | 140 | 38k | 0.783 | 0.623 | 0.694 |
| 7 | 1.5 | 95 | 54k (1.4×) | 0.783 | 0.614 | 0.688 |
| 6 | 2 | 71 | 70k (1.8×) | 0.785 | 0.596 | 0.678 |
| 5 | 2.5 | 57 | 84k (2.2×) | 0.787 | 0.585 | 0.671 |
| Species inclusion test |  |  |  |  |  |  |
| - | 1 | 140 | 47k | 0.9993 | 0.9873 | 0.9933 |
| 7 | 1.5 | 95 | 69k (1.5×) | 0.9989 | 0.9872 | 0.9930 |
| 6 | 2 | 71 | 90k (1.9×) | 0.9983 | 0.9871 | 0.9927 |
| 5 | 2.5 | 57 | 109k (2.3×) | 0.9981 | 0.9872 | 0.9927 |

Homology detection sensitivity decreased in the species exclusion test at higher compression rates. While syncmer selection effectively preserves the density of shared metamers, the lower absolute count of shared metamers makes distant homologs harder to detect. Despite this, the final configuration above with *s* = 5 achieved recall equivalent to the default Metabuli, with a 3-percentage-point increase in precision.

In the species inclusion test, higher compression rates resulted in more reads being classified at the species rank because syncmer subsampling breaks tie scores, reducing LCA rank elevations. This dynamic creates a slight trade-off: species-level recall increases as more assignments are made, but precision decreases slightly because some ties are resolved incorrectly.

### Comprehensive benchmarking of Metabuli in inclusion and exclusion tests

We compared various configurations of the improved Metabuli against both the previous Metabuli and other alignment-free taxonomic classifiers (Figure 2). To evaluate classification performance across diverse taxonomic ranks, we designed a series of exclusion tests. In the subspecies exclusion test, methods utilizing DNA-level searches (Kraken2 and Centrifuger) outperformed tools executing translated searches against a protein database. This advantage stems from the fact that nucleotide-level information provides the higher resolution necessary to distinguish between very closely related taxa. However, in the species exclusion test, accurate genus-level classification requires the detection of homology shared between different species within the same genus. In this scenario, protein-based software excelled because amino acid sequences are more highly conserved across these broader taxonomic distances. This same trend was consistently observed in the genus and family exclusion tests.

**Figure 2.**
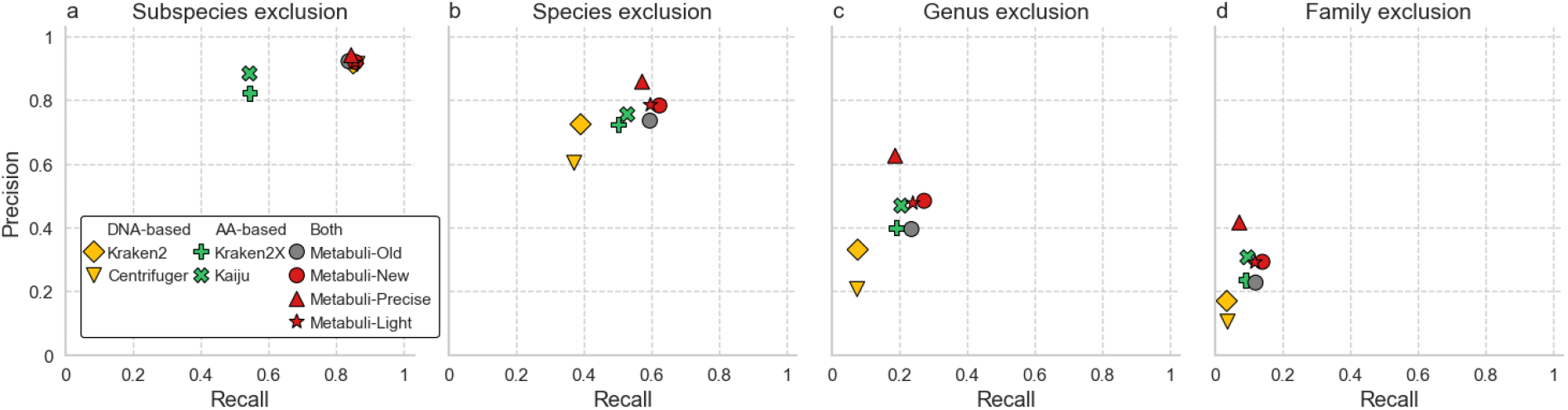
Taxonomic exclusion test results. Evaluation of homology detection sensitivity at the (a) species, (b) genus, (c) family, and (d) order levels. Accuracy was evaluated at the next highest rank of exclusion (species, genus, family, and order, respectively) because the query’s target clade was removed. Metabuli configurations: Old (v1.1.1), New (final configuration in Table 2), Precise (New + Precise preset (`-–min-score 0.2 --min-sp-score 0.6 -e 0.001)`), and Light (New + syncmer (*s*=6)). See Table S2 for actaul values.

**Figure 3.**
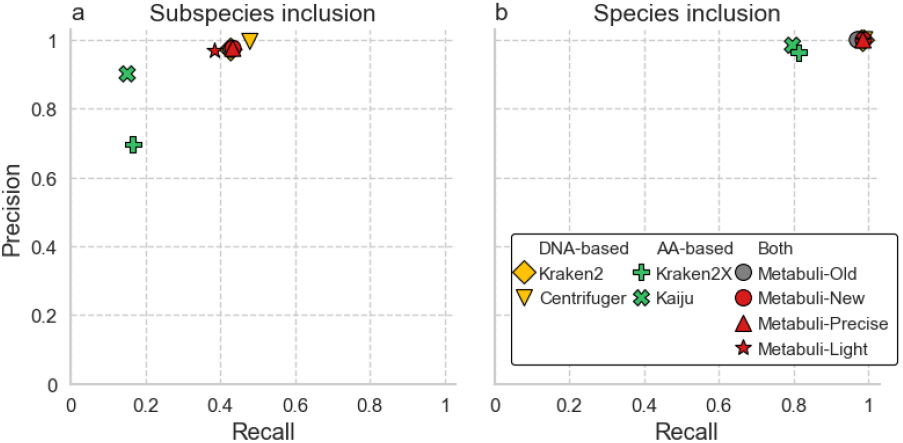
Taxonomic inclusion test results. Evaluation of classification resolution at the (a) subspecies and (b) species levels. The reference database contains at least one closely related sibling per queried taxon to challenge the algorithm’s classification resolution.

The previous version v1.1.1, Metabuli-Old, demonstrated performance comparable to DNA-based tools in the subspecies exclusion test, while matching the accuracy of protein-based tools in higher-rank exclusion tests. This versatility is achieved through Metabuli’s joint DNA and amino acid analysis using metamers (16). The new Metabuli configurations maintained this strong performance in the subspecies exclusion test, where sensitivity is less critical because the queried species remains in the database. However, in the higher-rank exclusion tests that demand sensitive homology detection, utilizing spaced metamers with a reduced amino acid alphabet (Metabuli-New) improved both precision and recall. Applying a minimum score threshold and E-value-based filtering to this configuration (Metabuli-Precise) significantly boosted precision. Furthermore, implementing closed syncmer sampling (Metabuli-Light) yielded a recall slightly lower than Metabuli-New, but still superior to the other baseline tools. Given its top-tier performance, halved database size, and doubled classification speed, Metabuli-Light is highly advantageous for processing massive datasets or operating in consumer-grade computing environments (17).

In the inclusion tests, Metabuli performed comparably to the DNA-based tools Kraken2 and Centrifuger, while outperforming the protein-based methods Kaiju and Kraken2X. Centrifuger achieved the highest resolution in distinguishing subspecies. This enhanced resolution likely results from its ability to identify flexible-length matches and its indexing strategy; unlike Kraken2, which collapses matches to the lowest common ancestor, or Metabuli, which collapses them at the species level, Centrifuger records the specific sequences that share a common substring. Metabuli-Light demonstrated reduced subspecies resolution, as syncmer sub-sampling naturally depletes the pool of subspecies-specific k-mers.

## Future Directions

In future studies, we aim to explore the utilization of consecutive joker positions within spacing masks (e.g., 1`1100111011`). Such patterns remain porous even when adjacent windows are merged, potentially enhancing mismatch tolerance. In the inclusion tests, despite encoding 27 nucleotides per metamer (an increase of 3 nucleotides over Metabuli-Old), Metabuli-New did not exhibit improved resolution at the species or subspecies levels. This suggests an opportunity to further optimize DNA utilization, balancing the algorithm so that gains in overall sensitivity do not confound high-resolution assignments.

Furthermore, we plan to extend our synthetic benchmarking to encompass long-read sequencing technologies, following the methodology of Kim et al. (16), and to evaluate the algorithm against independent datasets like CAMI2 (21) for standardized performance comparisons. Applying this advanced iteration of Metabuli to environmental metagenomics represents a highly relevant use case; these studies demand exceptional mapping sensitivity, as many environmental species are vastly underrepresented in public repositories.

## Conclusions

We improved the Metabuli classification framework by replacing its original metamer encoding with a more efficient bit-packing architecture. This structural update enabled the integration of reduced amino acid alphabets and spaced masks, which increased mismatch tolerance and improved homology detection for divergent sequences. Furthermore, integrating closed syncmer sub-sampling achieved a 50% reduction in database size and a twofold acceleration in processing speed while maintaining classification accuracy. In synthetic exclusion tests, Metabuli v1.2 showed higher classification sensitivity than v1.1.1 and other evaluated tools, while matching the specificity of DNA-based methods in inclusion tests. Altogether, these updates provide a scalable and effective solution for profiling complex environmental datasets.

## Supplementary Material

**Table S1.**
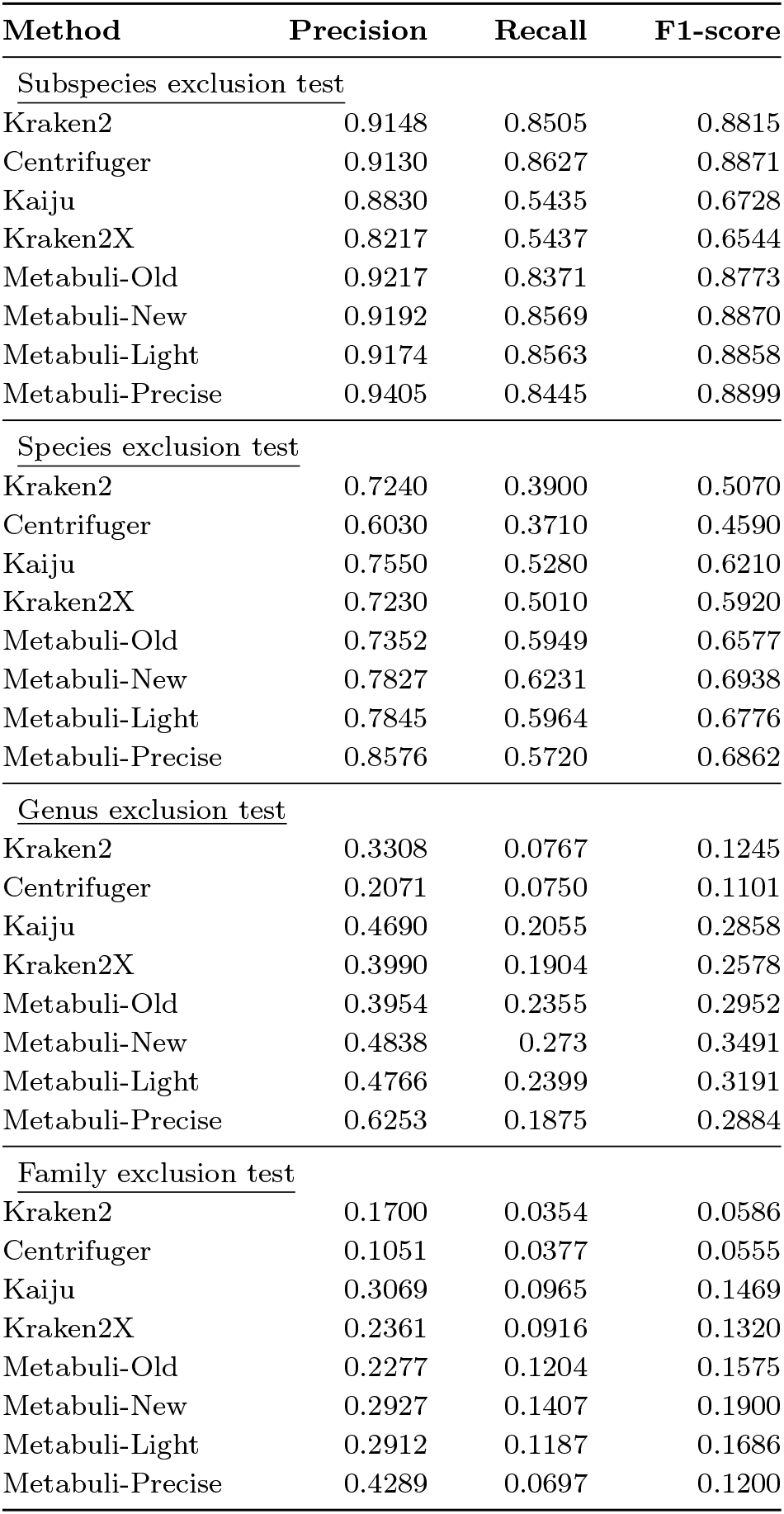
Taxonomic exclusion test performance metrics.

| Method | Precision | Recall | F1-score |
| --- | --- | --- | --- |
| Subspecies exclusion test |  |  |  |
| Kraken2 | 0.9148 | 0.8505 | 0.8815 |
| Centrifuger | 0.9130 | 0.8627 | 0.8871 |
| Kaiju | 0.8830 | 0.5435 | 0.6728 |
| Kraken2X | 0.8217 | 0.5437 | 0.6544 |
| Metabuli-Old | 0.9217 | 0.8371 | 0.8773 |
| Metabuli-New | 0.9192 | 0.8569 | 0.8870 |
| Metabuli-Light | 0.9174 | 0.8563 | 0.8858 |
| Metabuli-Precise | 0.9405 | 0.8445 | 0.8899 |
| Species exclusion test |  |  |  |
| Kraken2 | 0.7240 | 0.3900 | 0.5070 |
| Centrifuger | 0.6030 | 0.3710 | 0.4590 |
| Kaiju | 0.7550 | 0.5280 | 0.6210 |
| Kraken2X | 0.7230 | 0.5010 | 0.5920 |
| Metabuli-Old | 0.7352 | 0.5949 | 0.6577 |
| Metabuli-New | 0.7827 | 0.6231 | 0.6938 |
| Metabuli-Light | 0.7845 | 0.5964 | 0.6776 |
| Metabuli-Precise | 0.8576 | 0.5720 | 0.6862 |
| Genus exclusion test |  |  |  |
| Kraken2 | 0.3308 | 0.0767 | 0.1245 |
| Centrifuger | 0.2071 | 0.0750 | 0.1101 |
| Kaiju | 0.4690 | 0.2055 | 0.2858 |
| Kraken2X | 0.3990 | 0.1904 | 0.2578 |
| Metabuli-Old | 0.3954 | 0.2355 | 0.2952 |
| Metabuli-New | 0.4838 | 0.273 | 0.3491 |
| Metabuli-Light | 0.4766 | 0.2399 | 0.3191 |
| Metabuli-Precise | 0.6253 | 0.1875 | 0.2884 |
| Family exclusion test |  |  |  |
| Kraken2 | 0.1700 | 0.0354 | 0.0586 |
| Centrifuger | 0.1051 | 0.0377 | 0.0555 |
| Kaiju | 0.3069 | 0.0965 | 0.1469 |
| Kraken2X | 0.2361 | 0.0916 | 0.1320 |
| Metabuli-Old | 0.2277 | 0.1204 | 0.1575 |
| Metabuli-New | 0.2927 | 0.1407 | 0.1900 |
| Metabuli-Light | 0.2912 | 0.1187 | 0.1686 |
| Metabuli-Precise | 0.4289 | 0.0697 | 0.1200 |

**Table S2.** Taxonomic inclusion test performance metrics.

| Method | Precision | Recall | F1-score |
| --- | --- | --- | --- |
| Subspecies inclusion test |  |  |  |
| Kraken2 | 0.9716 | 0.4280 | 0.5943 |
| Centrifuger | 0.9949 | 0.4793 | 0.6469 |
| Kaiju | 0.9008 | 0.1511 | 0.2588 |
| Kraken2X | 0.6969 | 0.1656 | 0.2677 |
| Metabuli-Old | 0.9720 | 0.4291 | 0.5954 |
| Metabuli-New | 0.9749 | 0.4337 | 0.6003 |
| Metabuli-Light | 0.9673 | 0.3855 | 0.5513 |
| Metabuli-Precise | 0.9751 | 0.4333 | 0.6000 |
| Species inclusion test |  |  |  |
| Kraken2 | 0.9988 | 0.9860 | 0.9924 |
| Centrifuger | 0.9998 | 0.9904 | 0.9951 |
| Kaiju | 0.9838 | 0.7975 | 0.8809 |
| Kraken2X | 0.9632 | 0.8141 | 0.8824 |
| Metabuli-Old | 0.9997 | 0.9791 | 0.9893 |
| Metabuli-New | 0.9993 | 0.9873 | 0.9933 |
| Metabuli-Light | 0.9983 | 0.9871 | 0.9927 |
| Metabuli-Precise | 0.9993 | 0.9864 | 0.9928 |

